# Estradiol (E2) concentration shapes the chromatin binding landscape of the estrogen receptor

**DOI:** 10.1101/2022.09.23.509212

**Authors:** Amy L. Han, Kiran Vinod-Paul, Satyanarayan Rao, Heather M. Brechbuhl, Carol A. Sartorius, Srinivas Ramachandran, Peter Kabos

## Abstract

How transcription factors (TF) selectively occupy a minute subset of their binding sites from a sizeable pool of putative sites in large mammalian genomes remains an important unanswered question. In part, nucleosomes help by creating formidable barriers to TF binding. TF concentration itself plays a crucial role in the competition between TFs and nucleosomes. In the case of nuclear receptors, the ligand adds another layer of complexity. Estrogen receptor alpha (ER) is a classic example where its main ligand estradiol (E2) can modulate ER binding on chromatin. Here we show a complete rewiring of ER binding as a function of E2 concentration. As E2 concentration increases by two orders of magnitude, ER levels decrease, and ER binding localizes to promoter-distal sites with strong ER motifs. At low E2 levels, abundant levels of ER are present in the nucleus, and ER binding occurs mostly at sites without an identifiable ER binding motif, potentially in cooperation with other TFs like STAT1. We propose that E2’s effect on ER activity plays a major role in defining genome-wide ER binding profiles. Thus, variations in E2 concentrations in ER-positive breast tumors could be a significant factor driving heterogeneity in tumor phenotype, treatment response, and potentially drug resistance.

## Introduction

Breast cancer is the second leading cause of death from cancer in women in the United States (Siegel et al., 2022). About 75% of these breast cancer cases are estrogen receptor alpha positive (ER^+^). 17β-estradiol (E2) by binding and activating ER, plays a central role in breast cancer development and progression (Yager & Davidson, 2006). Anti-endocrine therapy remains the mainstay for treatment of ER^+^ breast cancer (Bundred et al., 2002; Fisher et al., 1998). Endocrine targeting is achieved either by modulation of ER activity by competitive inhibition, degradation/downregulation of the receptor itself, or use of aromatase inhibitors to reduce the local synthesis of E2. Unfortunately, patients’ response to these treatments vary (AlFakeeh & Brezden-Masley, 2018; Zelnak & O’Regan, 2015) and their effectiveness is limited by intrinsic (15-20%) and acquired endocrine resistance (30-40%) causing a major obstacle to successful therapy of the disease (Anurag et al., 2018; Lei et al., 2019). ER protein expression is also heterogeneous, and its expression does not always correlate with anti-endocrine treatment response (Kim et al., 2011). In addition, other factors like the E2 levels present in tissues or surrounding tumor microenvironment (TME) influence anti-endocrine treatment response. For instance, different fibroblast subtypes can influence breast cancer cells’ sensitivity to E2 and resistance to tamoxifen therapy (Brechbuhl et al., 2017). Thus, the impact of E2 concentration warrants further investigation, particularly lower E2 conditions typical of post-menopausal women and women on anti-endocrine therapy including ovarian suppression. Consideration of variable E2/ER activity may help identify potential prognostic markers and/or novel therapeutic targets.

Genome-wide analyses have revealed the mechanisms of E2/ER binding and signaling (Cheung & Kraus, 2010; Gilfillan et al., 2012) to be complex with E2-induction resulting in ER binding a wide range of distant sites including distal enhancers that lie within 50kb of ER-binding sites (Gertz et al., 2012; Hah et al., 2013; Lin et al., 2007). ER binding at these distal sites regulate transcriptional start sites (TSS) through the cooperation from other co-factors/partners like FOXA1, GATA3, and p300 (Farcas et al., 2021). Yet, we still do not understand how E2 modulates genome-wide ER binding profiles and how some of these cooperating factors affect the global ER transcriptome. The effect of E2 on ER binding site selection is especially important in comprehending anti-endocrine resistance and cancer progression in post-menopausal women. Even after acquiring anti-endocrine resistance, ER^+^ breast cancers tend to remain ER-dependent (Osborne & Schiff, 2011). ER “degraders” like fulvestrant emerged as a means to inhibit receptor dimerization through a ligand-induced conformation change that is targeted for proteasomal degradation (McDonnell & Wardell, 2010; Mottamal et al., 2021). However, recent studies propose suppression of ER intra-nuclear mobility by “ER degraders” rather than elimination of the receptor itself (Guan et al., 2019). Paradoxically, these E2-competitive ligands promote ER binding to chromatin, impacting their chromatin accessibility and thereby ER signaling response. This suggests both ligand type and concentration profoundly impact the ER cistrome and hence outcomes in breast cancer.

It is unclear how much E2 is present in ER^+^ breast tumors and how ER behaves based on its ligand availability, especially in the context of endocrine manipulation. Prior pre-clinical studies have used E2 concentrations ranging from 10^−14^ to 10^−7^ M, with the majority of studies using the higher end of E2 (10^−7^ M E2) when comparing to estrogen withdrawn conditions (EWD). In a study of E2 on the immunomodulatory role of immune cells, lower concentrations of E2 significantly increased the proliferation of splenic T lymphocytes and interferon gamma production (Priyanka et al., 2013). These E2 concentration dependent effects on cell-mediated immune responses indicate that even low E2 levels can exert significant biological effects. The current standard method of E2 measurement is by ELISA, which is limited in its sensitivity of detection (Rosner et al., 2013). In addition, tissue mass spectrometry measurement of E2 is not optimal nor routine due to difficulties in obtaining consistent results with minimal matrix effects and background interference (Denver et al., 2019; Keski-Rahkonen et al., 2013). Therefore, it is not well understood how much E2 is required for an active ER response. As an alternative to directly determining E2 concentrations in tissues, we sought to map the functional output of ER as a readout of E2 concentration. We mapped genome-wide ER binding across four orders of E2 concentration to build a ligand sensitive accessibility signature. We performed global analyses of ER binding across varying E2 concentrations using the highly specific <u>c</u>leavage <u>u</u>nder targets and release <u>u</u>sing <u>n</u>uclease (CUT&RUN) (Skene et al., 2018) in ER^+^ MCF7 and UCD12 breast cancer cell lines.

Our study addresses how various E2 concentrations modulate the ER binding program to prospectively understand what occurs under native E2 conditions present in ER^+^ breast cancer patients. We identified ER peaks that cluster according to E2 concentration. In higher E2 environments, the number of binding events associated with estrogen responsive elements (ERE) increases; however, the majority of the E2/ER cistrome does not involved EREs. This suggests that E2/ER has non-canonical motif binding or is being tethered through other factors. Specifically at lower E2 concentrations, indicative of a post-menopausal state, we observed an enrichment of non-canonical ER binding near STAT1 motifs. Knockdown of STAT1 decreased non-canonical ER binding, suggesting STAT1 involvement in ER binding at low E2 concentrations.

## Methods

### Cell Culture

Human MCF-7 (p53 wildtype, ER^+^) breast cancer cells were cultured in Dulbecco’s Modified Eagle Medium (DMEM) (Corning, 10-013-CV) supplemented with 10% Fetal Bovine Serum (FBS) (Atlas Biologicals, EF-0500-A). The ER^+^ UCD12 (Finlay-Schultz et al., 2020) cells were derived from patient-derived xenograft (PDX) and cultured in DMEM/Hams F-12 50:50 Mix (Corning, 10-092-CV) supplemented with 10% FBS. Cells were grown at 37^°^C with 5% CO_2_/95% atmospheric air. Breast cancer cell lines were placed in estrogen withdrawn (EWD) media: Minimum Essential Medium (MEM) no L-glutamine, no phenol red (Gibco, 51200-038) supplemented with 10% Dextran Coated Charcoal Stripped (DCC) FBS and glutagro supplement (Corning, 25-015-Cl) for 72 hours prior to use in experiments. All cell lines were authenticated by short tandem repeat profile testing. All cell lines were validated to be mycoplasma free before use in any experiments using the Universal Mycoplasma Detection Kit from American Type Culture Collection.

### Western Blot

Cells were collected 1 hour after E2 (10^−12^ M, 10^−10^ M, 10^−8^ M) or ethanol (ETOH) (Decon Labs, Inc.) treatment after being EWD for 72 hours. Cells were washed 2x’s with PBS and then lysed with NP-40 cell lysis (Invitrogen, FNN0021) buffer supplemented with Halt™ protease inhibitor cocktail, EDTA-free (Thermo Scientific, 78425), Complete phosphatase inhibitor (Roche, 04693132001), and Phenylmethylsulfonyl Fluoride (PMSF) (Thermo Scientific, 36978). Protein quantification was done using Pierce™ Detergent Compatible Bradford assay (Thermo Scientific, #1863028). 30mg of protein with the addition of NuPAGE® LDS sample buffer (4x) (Invitrogen, NP0007) + NuPAGE sample reducing agent (10x) (Invitrogen, NP0009) were added to the wells and ran on a NuPage™ 4-12% Bis-Tris mini gel (Invitrogen, NP0336BOX). After gel electrophoresis, protein was transferred to a PVDF membrane (Millipore, IPFL00010). The blot was blocked for 1 hour in LI-COR Odyssey Blocking Buffer PBS (LI-COR, 927-40000), then incubated with primary antibody overnight gently rocking in 4°C. The membrane was then washed 3x’s with TBST followed by appropriate secondary LI-COR antibody. Images of the protein expression were detected and taken with the LI-COR Odyssey system.

### qRT-PCR

Cells were washed 2x’s with Phosphate Buffered Saline (PBS) (Corning, 21-040-CV) and collected 1 hour after E2 (10^−12^ M, 10^−10^ M, 10^−8^ M) or ETOH treatment after 72 hours of EWD. RNA was extracted using the Direct-zol RNA Miniprep (Zymo Research, R2052) and cDNA was synthesized using SuperScript III First-Strand Synthesis System (Invitrogen, 18080-051). qPCR was performed using PerfeCTa SYBR Green SuperMix (QuantaBio, 95054) on the BioRad CFx96 Touch Real-Time PCR Detection System. Relative transcriptional expression was quantified and normalized.

### CUT&RUN

After 72 hours of EWD, cells were plated in 100mm dishes for CUT&RUN. Cells were either induced with 10^−12^ M, 10^−10^ M, 10^−8^ M of E2 or ETOH for one hour prior to being washed twice with PBS and harvested using a cell lifter. Cells were counted to have 5 × 10^5^ cells per sample. The protocol was conducted as specified in Skene P et al. (Skene et al., 2018) in triplicates for each sample condition. Briefly, concanavalin-A beads (Bang Labs Inc.) were washed and activated with Binding Buffer, then added to the cells for 10 minutes at room temperature (RT) while rotating. The buffer was removed after being cleared on the magnetic stand in which the bound cells to the activated beads were resuspended in Antibody Buffer (100ml/sample). 1ml of respective ER or IgG antibody was added to each sample tube and incubated overnight at 4°C. Samples were washed with 0.025% Digitonin Buffer three times before adding in pA-MNase (EpiCypher, R&D151016). Tubes were placed for 1 hour at 4°C while nutating to allow the binding of pA-MNase. Samples were washed thoroughly with the Digitonin Buffer three times, resuspended in Digitonin Buffer, and placed on a pre-chilled heating block on wet ice. 2ml of 100 mM CaCl_2_ was added to each tube with gentle vortexing and immediately placed back into the 0°C block. Tubes were then incubated at 4°C for 2 hours with nutation for targeted chromatin digestion. Reaction was stopped with the addition of 100ml of STOP Buffer and mixed by gentle vortexing. Samples were incubated at 37°C for 30 minutes on a ThermoMixer at 500rpm to release the fragments into solution. Tubes were placed on a magnet stand and the supernatant was transferred to a new Eppendorf tube. DNA extraction followed using the phenol-chloroform-isoamyl alcohol 25:24:1 (PCI; Invitrogen, 15593049) method as described in Skene P et al. (Skene et al., 2018).

### CUT&RUN libraries

CUT&RUN libraries were made using the Ovation Ultralow System V2 kit (Tecan, 0344NB-A01) with the specified PCR conditions for library amplification: 98ºC for 45s, (98ºC for 15s, 60ºC for 10s) x 13 cycles, 72ºC for 1 min, followed by 4ºC hold and library purification with AMPure XP beads (Beckman Coulter, A63881). End repair, ligation, and ligation purification were done as specified by the kit’s specified protocol. After DNA TapeStation analysis (Agilent), libraries were submitted to the Genomics and Microarray Shared Resource at the University of Colorado Anschutz Medical Campus for sequencing of 10 million paired end reads per sample on the NovaSEQ6000 (Illumina).

### CUT&RUN Data Processing and Peak Calling

CUT&RUN libraries were aligned to the human (GRCh38) using Bowtie2 v2.3. The aligned files were converted to BEDPE/BED files using bedtools v2.18. Then using custom python scripts and *wigtobigwig()* from UCSC tools (https://hgdownload.cse.ucsc.edu/admin/exe/linux.x86_64/), the BED files were converted into a spike-in normalized coverage wig/bigwig files and only the short fragments with length 30-121bp were considered. For peak calling, custom python script where normalized coverage of the short fragments (at 10bp resolution) was smoothened using Savitzky-Golay filter, which is available as SciPy function ‘signal.savgol_filter()’ with parameters window_length = 9, polyorder= 1. The cutoff for each dataset was determined by iteratively eliminating outliers and used ‘find_peaks()’ function in SciPy call peaks that are separated by a minimum of 250 bp.

### Differential Peaks

For each concentration, ETOH and IgG condition CUT&RUN scores were calculated as the read density in regions spanning the CUT&RUN peak summit ± 30bp. These scores were used to calculate differential peaks among two conditions where the log2 ratio of the two conditions were computed and if the log2 fold change is >1 or <-1 the peaks are grouped in condition1 and 2 and the remaining peaks were grouped as common. Based on these groups the profile plots are plotted where the ±1000 bp flanking regions were considered from the peak summit and the normalized read density was calculated in each base pair for ETOH and E2 conditions. The Venn diagrams from the differential groups of the three concentrations were generated using EVenn (http://www.ehbio.com/test/venn/#/). The peak intersection was done by bedtools v2.27.1.

### Peak Clustering

The differential E2 peaks from all the concentrations were combined and k-means clustering was applied on the Z-scores of the peaks, using the function kmeans() from factoextra r package. 6 clusters were found out of which one cluster was discarded from the further analysis due to its mixed definitions in all the conditions from the rest of the clusters. Using the 5 defined cluster definitions, the CUT&RUN score (which is determined by the density of the reads) was generated and a heatmap was generated using the R package, ComplexHeatmap.

### Motif Discovery and Fractions

Motif discovery was done in the peaks from each cluster using command line version of FIMO from MEME-SUITE. All human motifs from HOCOMOCO v11 database were used. Transcriptions factors (TF) that are expressed (normalized counts >= 5) were considered for further analysis. Following motif discovery, the transcription factor (TF) fraction in each cluster was calculated. To see the pattern of the fractions from cluster 1 to cluster 5, each TF fraction from all the clusters were then clustered by Kmeans() from amap R package and correlated using Spearman Correlation.

### STAT1 Knockdown

STAT1 human siRNA Oligo Duplex and nontargeting scramble siRNA (Origene) were transiently transfected in the MCF7 cells. Briefly, cells were plated in EWD media for 48 hours, changed with fresh EWD media prior to transfection with 50nM of respective siRNA using siTran2.0 reagent (Origene). Fresh EWD media was replaced 18 hours post-transfection and cells were left in the 37^°^C incubator for an additional 48 hours. Transfected cells were treated with either 10^−12^ M E2 or 10^−8^ M E2 an hour before harvesting the cells for ER CUT&RUN assay or protein lysis for western blot analysis of STAT1 expression.

### Immunoprecipitation

72 hours after EWD, cells were treated with either 10^−12^ M or 10^−8^ M of E2 or ETOH for one hour prior to being washed twice with PBS and harvested using a cell lifter. Cells were collected in PBS, centrifuged, and PBS was aspirated. Cold nondenaturing lysis buffer + HALT protease inhibitor (Thermo Scientific) was added and mixed thoroughly to obtain protein lysates. Lysates were pre-cleared with Protein A dynabeads (Invitrogen, 10002D) + IgG under rotary agitation for 45 min at 4ºC. IgG, ER, or STAT1 antibody was bound to the Protein A dynabeads by incubation of the antibody bead mixture for 2 hours at 4ºC by gentle rotary mixing. Antibody-beads were crosslinked with the addition of 5mM bis(sulfosuccinaimidyl)suberate (BS3) for 30 minutes by gently mixing. Crosslinking reaction was quenched by addition of 1M Tris-HCl pH7.4 to each sample with incubation at RT under rotary agitation for 15 minutes. Cross-linked beads were washed once with 1x PBS, followed by 2x’s wash step with wash buffer. 500mg of cell lysates were added to the sample tubes in 300ml total volume. Lysates and bead-antibody conjugate mixture were incubated at 4°C overnight under rotary agitation. Samples were washed 3x’s with wash buffer and eluted with 1x loading buffer (1% SDS/0.5% NP-40) and incubated for 10 minutes at 100°C. ERα IP supernatants were collected and run on ProteinSimple-Jess.

## Results

### Impact of E2 concentration gradient on ER activation and cycling in breast cancer cells

To identify changes in ER binding as a function of E2 concentrations, we performed ER CUT&RUN on ER^+^ MCF7 and UCD12 cell lines (Fig. 1A). The UCD12 cell line was established from a patient with early-stage ER^+^ disease and is well characterized (Finlay-Schultz et al., 2020). Cells underwent EWD for at least 72 hours and then treated with varying concentrations of E2: 100-fold incremental doses starting from 10^−12^ M E2 and ending at 10^−8^ M E2 to encompass the presumed tissue concentration of E2 in breast cancer. After one hour of treatment, we washed and collected the cells for CUT&RUN.

**Figure 1.**
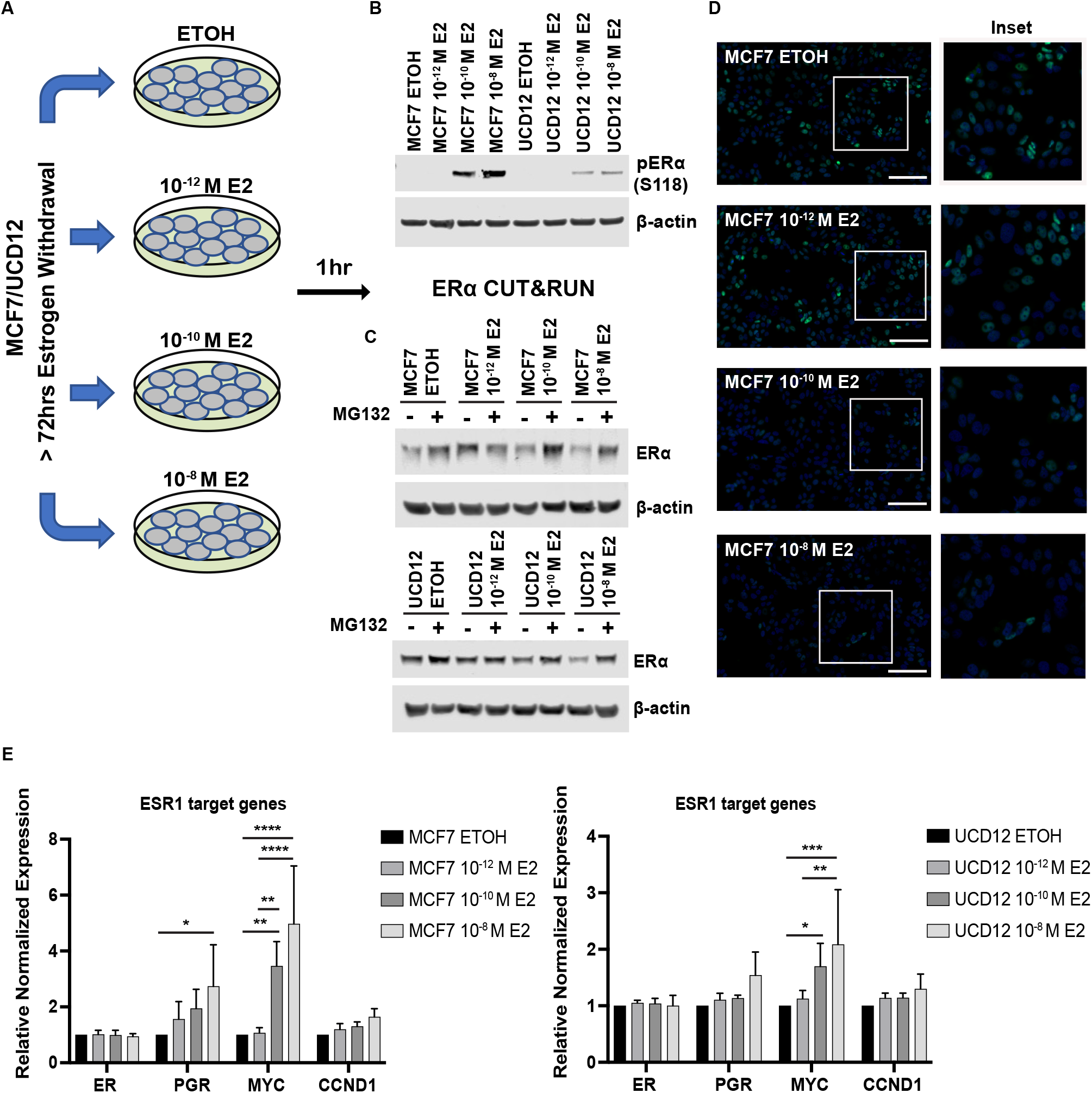
Graphical design and validation of E2 concentration effects on ER. A) Schematic representation of the experimental setup for the E2 concentration ER CUT&RUN. After EWD for >72 hours, MCF7/UCD12 cells were induced with estradiol (E2) across four orders of magnitude for one hour prior to being collected for downstream experiments (i.e., immunoblot, immunofluorescence, ER CUT&RUN). B) Western blot results for phospho-S118 ER (pERα) expression (pER. β-actin used as loading control. C) Immunoblot results for ERα protein expression with and without MG132, proteasome inhibitor. Cells were treated with either 10 μM MG132 or DMSO for 5 hours prior to being treated with E2 or ETOH (negative control) for 1 hour, then collected for immunoblot analysis. D) Representative immunofluorescence images of MCF7 cells 1 hour after addition of ETOH or different E2 concentrations with ER antibody (green) and DAPI (blue). 20X images, scale bar indicates 100 μm. Inset made from the boxed outline represents 200% zoomed in image. E) qPCR expression of ER and ESR1 target genes after 1 hour of ETOH/E2 treatment of EWD MCF7 and UCD12 cells with standard deviation. 2-way ANOVA statistical analysis, *p ≤ 0.05, **p ≤ 0.01, *** p ≤ 0.001, **** p ≤ 0.0001.

First, to confirm the validity of our experimental system, we examined the effect of different E2 concentrations on ER / phospho-ER protein levels in the MCF7 and UCD12 cells. Within an hour of E2 treatment, ER activation measured by phopho-S118 ER was observed starting at 10^−10^ M E2 (Fig. 1B). The increase in phospho-S118 was noticeably higher at the 10^−8^ M E2 concentration. Higher E2 concentrations also accompanied a decrease in total ER protein levels, as expected given the ligand-dependent feedback loop that triggers ER degradation. Treatment with proteasomal inhibitor MG132 in both MCF7 and UCD12 cells rescued the E2-dependent reduction in ER levels (Fig. 1C), suggesting the observed ligand-dependent ER recycling is due to the proteasomal degradation pathway as previously reported (Nawaz et al., 1999). Immunofluorescence staining also showed fewer ER^+^ cells with higher E2 concentrations (Fig. 1C, Fig. S1). Nuclear ER staining, including in ETOH control cells, confirmed the predominantly nuclear localization of ER both in the absence and presence of ligand. Overall, nuclear ER levels were highest in low E2 conditions and E2-activation of ER coincided with a rapid decrease in global nuclear ER levels.

We next examined transcriptional activation of select genes upon addition of E2. At the transcript level, ER target genes, PGR, MYC, and CCND1 increased with increasing E2 concentrations (Fig. 1D). MYC expression significantly increased in both MCF7 and UCD12 cells with higher E2, whereas PGR expression significantly differed between ETOH and 10^−8^ M E2 conditions in only the MCF7 cells. Collectively, these data demonstrated that ER activation and protein turnover and transcript levels of ER target genes are impacted by E2 abundance, establishing this system as robust for studying ER chromatin binding.

### E2-specific peaks increase non-linearly with E2 concentration

We evaluated the impact of the E2 gradient on ER binding from the CUT&RUN experiments at representative genomic locations. In line with increased PGR transcript expression, we also observed increased ER binding at the reported ER distal (+70,633 bp) binding site downstream of PGR (Palaniappan et al., 2019) at higher E2 concentrations (Fig. 2A). We then generated metaplots of ER CUT&RUN at peaks identified at each E2 concentration. We observed a gradual increase in CUT&RUN signal for E2-specific peaks with increased E2 concentrations in MCF7 (Fig. 2B) and UCD12 cells (Fig. S2). We calculated CUT&RUN scores for each peak and plotted the distribution of these scores as boxplots, which reiterated our observation with metaplots: an increase in the score of E2-specific peaks with increasing E2 concentration (Fig. 2C). We then identified three sets of peaks for each E2 concentration when compared with no E2 condition (ETOH): i) peaks that were common between the given E2 concentration and ETOH, ii) peaks that were unique to that concentration compared to ETOH, and iii) peaks that were seen only in ETOH in comparison to the given E2 concentration. Finally, we compared these three sets of peaks between the different E2 concentrations using Venn diagrams (Fig. 2D). The peaks that were common between any given E2 concentration and ETOH and the peaks that were specific to ETOH in comparison to any given E2 concentration featured significant overlaps among the three E2 concentrations. Strikingly, the peaks that were E2-specific at each concentration did not overlap between the different E2 concentrations. Thus, there was robust ER binding at all E2 concentrations and the E2-driven cistrome is unique to each E2 concentration.

**Figure 2.**
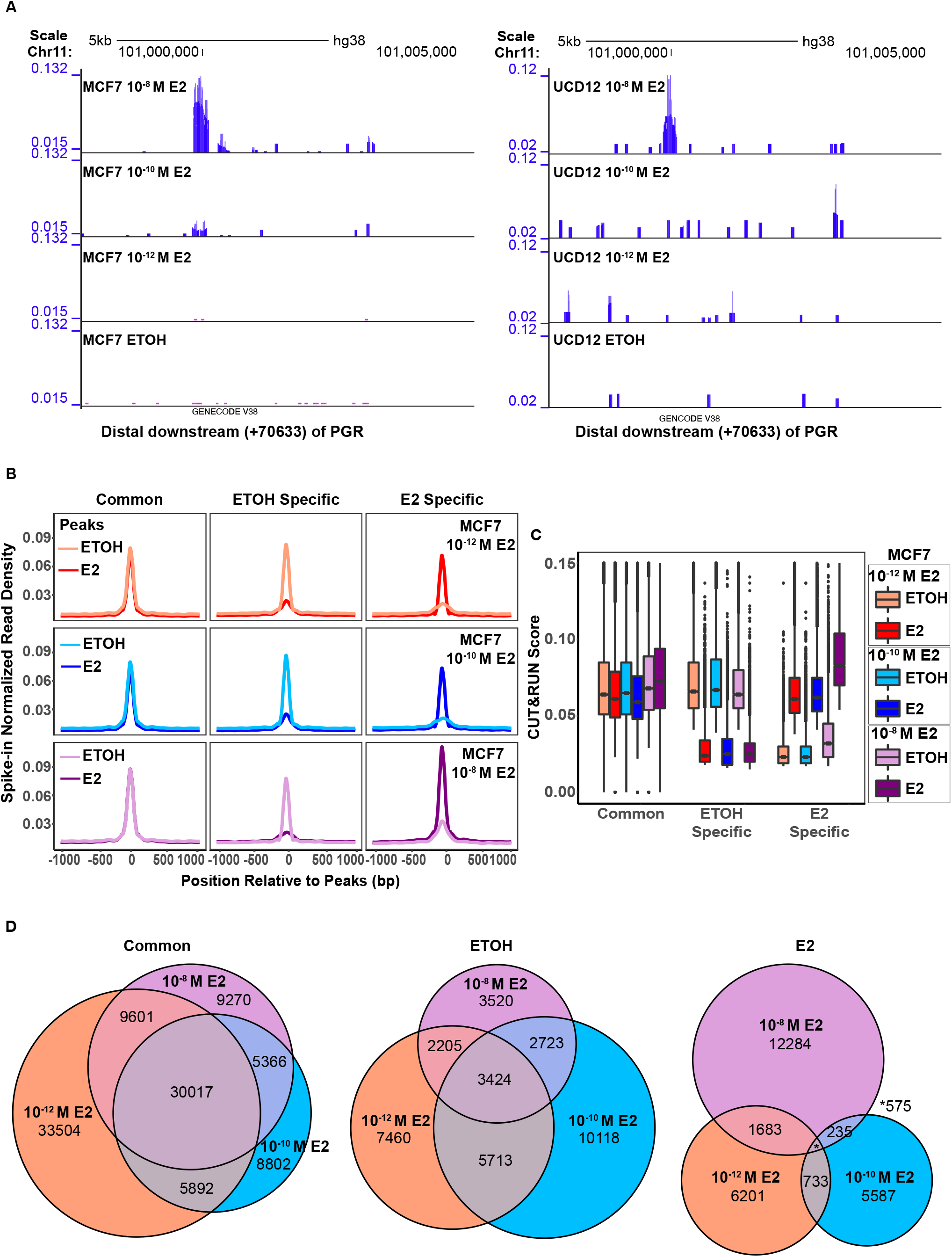
Patterns of ER binding based on E2 concentration. A) Track of ER CUT&RUN peaks +70633 bp distal downstream of PGR generated using UCSC Genome Browser. B) Profile plots from the differential peaks generated from MCF7 data. 1000 bp +/-as flanking regions from the peak summit and the spike-in normalized read density was calculated and plotted. C) Boxplot showing the CUT&RUN score distribution of the different E2/ETOH groups. D) MCF7 Venn diagrams representing the common and unique differential peaks among the common, ETOH-specific, and E2-specific groups between the three concentrations.

### Differential ER cistromes across E2 concentrations, predominantly involve non-canonical ER binding sites

To dissect concentration-specific changes in ER binding, we first created a super-set of peaks across datasets of CUT&RUN at different E2 concentrations, followed by k-means clustering of the spike-in normalized CUT&RUN scores at these peaks, with k=5. Visualizing CUT&RUN scores as a heatmap, ordered by clusters, IgG scores were the lowest, confirming the specificity of the ER CUT&RUN peaks (Fig. 3A, B). Under ETOH treatment, only a small set of peaks (cluster 5) displayed significant CUT&RUN scores, further confirming the capture of E2-dependent ER binding (Fig. 3A, B). Furthermore, the clusters resolved concentration-specific ER peaks into two main categories: The first category consisted of peaks specific to single E2 concentrations and include Clusters 1-3. Cluster 1 represented peaks that show significant binding only at 10^−12^ M E2, with Clusters 2 and 3 containing peaks with significant binding only in 10^−10^ M and 10^−8^ M E2, respectively. The second category comprised of peaks in Clusters 4 and 5, which displayed a progressive increase in E2-dependent CUT&RUN scores in a manner that would be expected for ligand-dependent activation. Ligand-dependent activation was further supported by the fact that these clusters had the highest proportion of peaks with canonical ERE motifs (Fig. 3C). In general, most of the E2-specific peaks were found distal (>1000bp) to the TSS, but the same trend favoring ERE peaks held true regardless of distance. The cluster with the next highest proportion of peaks with ERE motifs was the high-E2 specific peak cluster (cluster 3; Fig. 3C, Table S1), which also had the highest number of E2-specific differential peaks (4651). Interestingly, approximately 20% of these peaks had EREs, suggesting that even with high E2 ligand, the E2-ER response does not principally involve classical ERE binding. Thus, a significant fraction of ER binding did not involve sites with ERE motifs indicating the presence of alternative cofactors which might assist ER to bind DNA.

**Figure 3.**
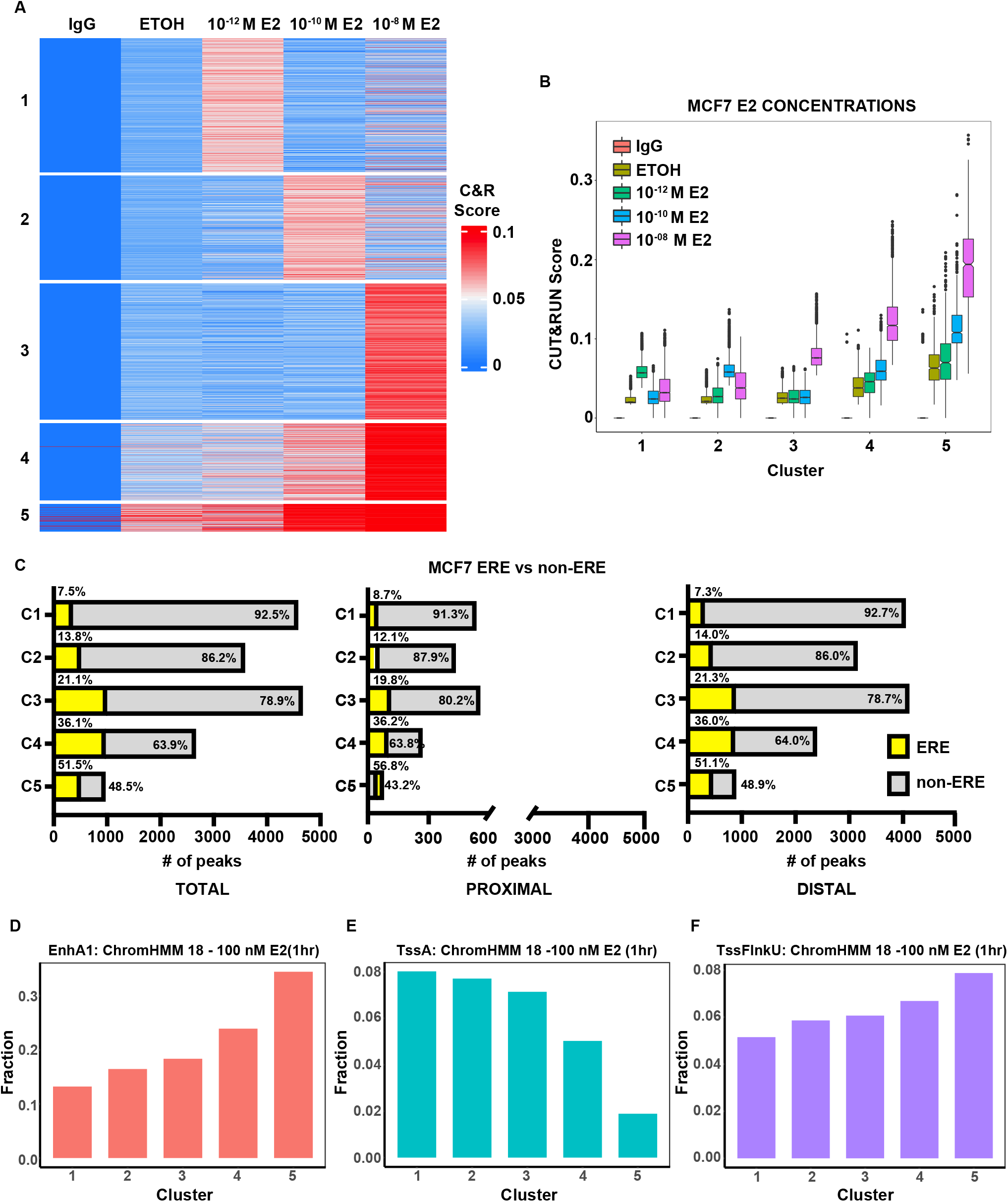
Differential ER binding in E2-specific peaks. A) Heatmap based on k-means clustering of the CUT&RUN scores at the E2-specific super-set of peaks from the different E2 concentrations, with k=5. B) CUT&RUN score distribution of the differential E2 binding clusters represented in boxplots. C) Graphs representing the number of peaks with ERE (yellow) or non-ERE (gray) and their percentages found in the total, proximal, and distal regions of ER binding. D-F) ChromHMM-18 states of the 100 nM/10^−7^ M E2 1 hour treated MCF7 samples. Graphs (based on fraction) of the groups that displayed a difference between the clusters. (D) EnhA1: Active Enhancer 1; (E) TssA: Active Transcriptional Start Sites; (F) TssFlnkU: Upstream Flanking TSS.

To determine how the peaks in each cluster were distributed across different parts of the genome, we intersected the peaks with the ChromHMM-18 annotation generated for 1 hour 100 nM E2 treatment and the two untreated samples by the Encode project (https://www.encodeproject.org/annotations/ENCSR151CZJ/). Out of the 18 ChromHMM-18 states (Fig. S3), three states showed a trend across the CUT&RUN clusters (Fig. 3D-F). Active enhancer (EnhA) state was lowest in cluster 1 and changed monotonically until cluster 5 (Fig. 3D). This was also true for the flanking TSS upstream (TssFlnkU) albeit to a much smaller degree (Fig. 3F). We saw a reverse trend for active TSS (TssA) activity, indicative of active promoters, with cluster 1 having the highest fraction of peaks with this state and cluster 5 having the lowest (Fig. 3E). These analyses showed that cluster 1, 2, & 3 binding sites are located in distinct parts of the genome in comparison to cluster 4 & 5. Our analysis in the absence of E2 and across E2 concentrations not only showed robust activation of ER binding by E2 at canonical ER motifs, but also widespread binding to sites lacking canonical ER motifs, which was heightened at lower E2 levels.

### Other TFs drive ER binding in low E2

A distinct set of sites bound by ER at low E2 concentrations, especially with a low fraction of ER motifs led us to hypothesize that binding in low E2 environments could be driven by ER-associated factors. ER typically functions as part of a complex made up by many factors. Included in this complex are multiple co-activators and co-repressors regulating the E2-driven transcriptional program (Farcas et al., 2021). To ask what other factors might be primarily driving ER binding in low E2 conditions, we screened all ER CUT&RUN peaks for known human TF motifs using FIMO. We then calculated the fraction of peaks with each motif within the E2 peak clusters and then filtered for TFs expressed in MCF7 cells in at least one of the E2/ETOH conditions that we employed. When we clustered motif enrichment across E2 groups, we found two predominant patterns (Figure 4A): Pattern 1/cluster 1 (C1) represented motifs enriched in low E2 peaks relative to high E2 peaks and pattern 2/cluster 2 (C2) represented motifs enriched in high relative to low E2 peaks.

**Figure 4.**
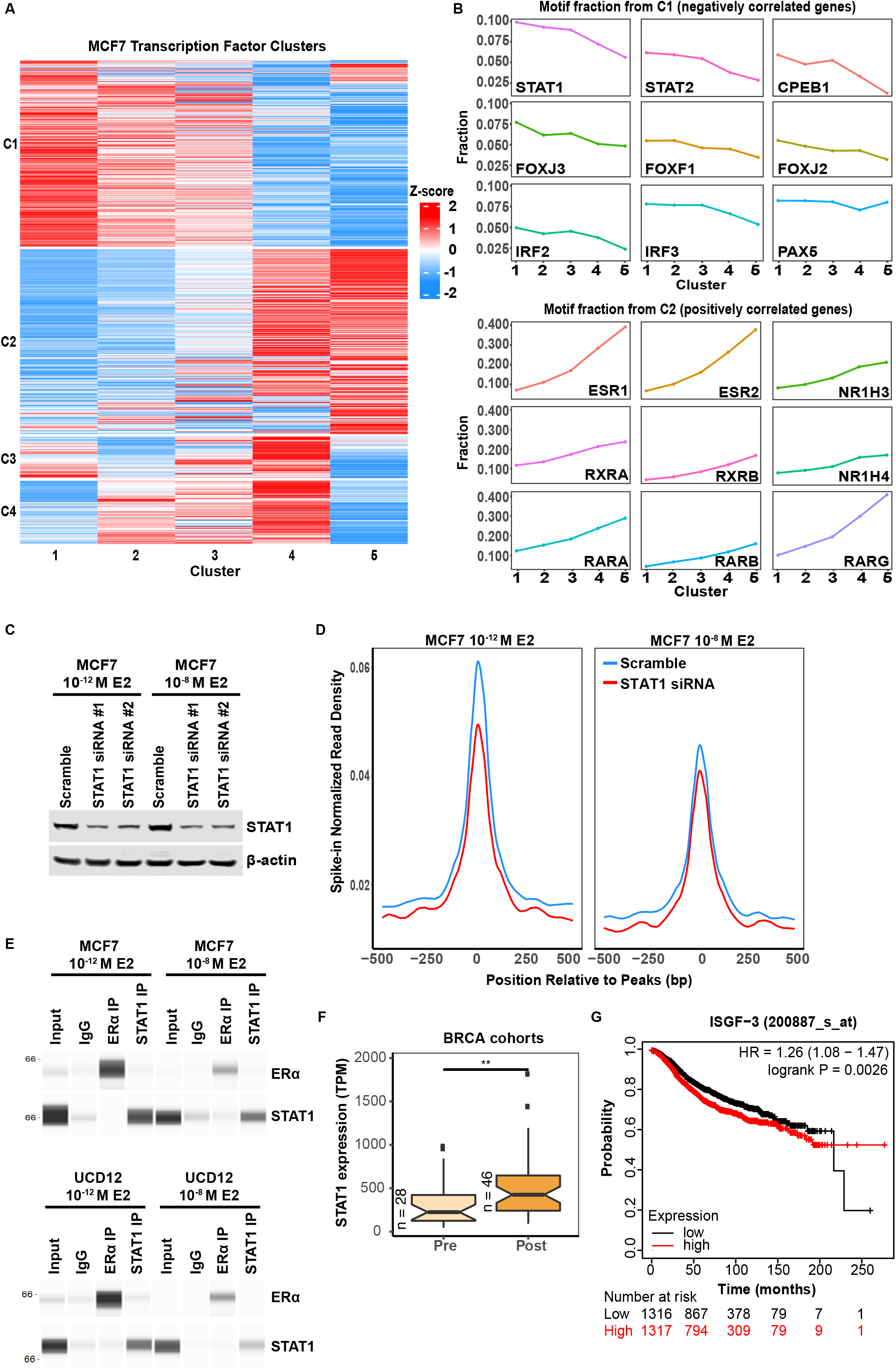
STAT1 influences ER binding at low 10^−12^ M E2 concentration. A) Heatmap based on the K means clustering of TF fractions in each CUT&RUN cluster. B) Top negatively and positively correlated TF identified from FIMO motif discovery. C) Western blot analysis of knockdown STAT1 protein expression from the STAT1 siRNA #1 and siRNA #2 vs scramble control transfection. D) Profile of the spike-in normalized read density of the STAT1 knockdown cells in the 10^−12^ M E2 and 10^−8^M E2 against the scramble control. E) Immunoprecipitation of ER and STAT1 with the 10^−12^ M E2 and 10^−8^M E2 induced MCF7 and UCD12 cells. F) TCGA data of STAT1 expression based on pre- and post-menopausal patients in the breast cancer cohort. G) Kaplan-Meier survival plot for STAT1 expression in ER^+^ breast cancer patients.

ER (ESR1 and ESR2) motifs were part of C2 and had the highest proportional increase in motifs in the clusters from the 10^−12^ M E2 group to the 10^−8^ M E2 as expected based on our analysis in Fig. 3. A group of other nuclear receptors, nuclear receptor subfamily 1 group H member (NR1H), retinoid x receptor (RXR), and retinoic acid receptor (RAR) TFs were also identified as positively correlated with the increase in E2 concentrations. RAR, an estrogen target gene, has been reported to co-occupy regulatory regions together with ER when cells are exposed to E2 (Ross-Innes et al., 2010). This binding occurred only at ERE sites and greatly diminished with ERE half-sites in an ER-dependent manner. RXR forms heterodimers with numerous other nuclear receptors and can bind multiple hormone response elements including EREs (Segars et al., 1993). RXR’s regulatory role is E2 dose-dependent and element-specific, only occurring in the presence of E2-activated ER. Thus, the TFs identified in C2 are supported by previous reports on other nuclear receptors associated with E2 induction, which supported our ability to discover new regulators involved in E2-dependent ER binding through this approach.

### STAT1 involvement in ER binding at the lower 10^−12^M E2 concentration

Based on us recapitulating known ER biology with top motifs in C2, we next asked which motifs drove ER binding at low compared to high E2. In C1, STAT1 was the top motif and it decreased in occurrence with increasing E2 concentrations. STAT2 also showed a negative correlation albeit to a lesser degree compared to STAT1. Other top motifs in C1 included cytoplasmic polyadenylation element binding protein 1 (CPEB1), B-cell-specific activator protein (PAX5), family of forkhead box (FOX) transcription factors as well as interferon-regulatory factors (IRFs).

To ask if the TFs binding these motifs promoted ER binding to DNA at low E2, we decided to focus on STAT1 as a representative TF. If STAT1 promoted ER binding at low E2 concentrations, we hypothesized that the loss of STAT1 would decrease ER binding. Hence, we performed ER CUT&RUN after knocking down STAT1. We reduced STAT1 levels with two different siRNAs (Fig. 4C) and then treated the cells with either 10^−12^ M E2 or 10^−8^ M E2 followed by ER CUT&RUN. Strikingly, knockdown of STAT1 resulted in significantly reduced CUT&RUN enrichment at 10^−12^ M E2, but only a minor reduction at 10^−8^ M E2 (Fig. 4D), suggesting STAT1 plays a role in ER binding that is specific to low E2 concentration. To determine if there was direct interaction between STAT1 and ER, we performed co-immunoprecipitation. Our results did not indicate direct binding between ER and STAT1 pulldown experiments, suggesting the STAT1 influence on ER binding was not due to direct interaction between the two TFs (Fig. 4E). In summary, STAT1 influenced ER binding at low E2 concentration, possibly indirectly by creating accessibility at non-canonical ER binding sites.

### STAT1 expression correlates with survival in ER^+^ breast cancer

Since we observed that STAT1 influences ER cistrome at low E2 concentrations, we asked if it might play a role in breast cancer in postmenopausal women, where E2 concentration is expected to be low. We observed higher STAT1 expression in postmenopausal compared to premenopausal women in TCGA cohorts (Fig. 4F). Furthermore, women with ER^+^ breast cancer and high STAT1 expression showed lower survival compared to women with low STAT1 expression that was statistically significant (Fig. 4G). This suggests STAT1 is a prognostic factor for lower survival, prospectively by regulating ER binding in low E2/postmenopausal conditions.

## Discussion

In this study, we found that the ER binding signatures at different E2 concentrations are unique. Our results reveal the diverse ways ER binds to chromatin in response to the availability of its ligand E2, which in breast cancer patients is a function of both menopausal status and therapy. Plasma E2 ranges from 30-400 pg/ml in premenopausal and 0-30 pg/ml for postmenopausal women; however, we do not have reliable E2 measurement in tissues to date (Rosner et al., 2013; Yaghjyan & Colditz, 2011). Therefore, little is known about the local estrogen levels in normal breast tissue or breast cancer (Yaghjyan & Colditz, 2011). We sought to address this issue by conducting ER CUT&RUN to detect ER binding activity across a range of E2 concentrations to serve as a potential surrogate for identifying E2 levels in tissue. Our assays were done an hour after treatment to identify the immediate effects of E2 induction and response. ER promoter occupancy has been reported to peak at 45 minutes after E2 treatment at higher, 10^−7^ M E2 concentrations (Shang et al., 2000). With the lower E2 amounts, we used a timepoint to capture ER binding induced even at 10^−12^ M E2. We recognize longer treatment times may be necessary to identify long-term effects. However, even with one hour of E2 treatment, we detect ER activation as indicated by the presence of phospho-ER starting at 10^−10^ M E2 in two ER^+^ breast cancer cell lines. Total ER expression followed a well-described pattern of degradation with increasing E2 concentration. These findings are consistent with reports demonstrating rapid degradation of ER protein in an E2-dependent manner (Alarid et al., 1999; Nawaz et al., 1999) and confirmed by inhibiting the proteasome with MG132, which prevented ER degradation in presence of E2.

Although we did not observe phospho-ER expression with 10^−12^ M E2, transcriptional expression of ER target genes shows a slight increase even at 10^−12^ M E2 compared to ETOH control. A similar trend was observed in the ER CUT&RUN enrichment at the known ER distal downstream region of PGR: Peak intensity present at higher E2 concentrations accompanies higher PGR transcriptional expression. With the increase in E2 concentration, we see a similar increase in E2 spike-in normalized read density. Interestingly, this incremental difference is more distinct with UCD12 cells. UCD12 cells have more heterogeneous ER expression compared to MCF7 cells, with reduced number of overall peaks obtained from the CUT&RUN (∼30,000 average number of peaks) compared to the MCF7 cells (∼50,000 average number of peaks). Nevertheless, ER CUT&RUN with both cell lines show similar results revealing the reproducibility of the assay and confirming the biological effects of E2 treatment. The E2-specific peaks do not display much overlap amongst the three concentrations, especially in comparison to the common and ETOH peaks, indicating that availability of the ligand, E2 dictates different ER binding that is not indicative of a linear response.

ER binding often occurs in regions distant to transcribed gene promoters. Lin et al. (Lin et al., 2007) reported that the majority of ER binding sites are located in either introns or are far distant to coding regions of genes with 43% of genes lying within 50 kb to an ER-binding site. Like with other genome-wide ER binding studies (Carroll et al., 2006; Kolendowski et al., 2018), we also find that most of the E2/ER binding does not occur at EREs, highlighting the cooperation of other factors and diverse functionality of ER in mediating transcriptional activity.

STAT1, primarily known for its involvement with interferon signaling, is a TF that plays a role in the transduction of stress and cytokine response, DNA damage, and activation of immune cell responses (Bailey et al., 2012). In cancer, STAT1 appears to have a dual role in its ability to be either oncogenic or tumor suppressive, which is dependent on cell type or environment. In our studies, STAT1 was the top motif found in our FIMO discovery under the lowest E2 condition, 10^−12^ M. Knockdown of STAT1 leads to the attenuation of ER CUT&RUN enriched peaks, which is only apparent with the lower 10^−12^ M E2 concentration. This binding mode of ER would be significant in ER^+^ tumors of post-menopausal women, where E2 levels are usually low. Our data does not support direct interaction between the two TFs; however, this does not eliminate the possibility of ER and STAT1 working together and being present in the same complex. For instance, STAT1 is known to bind to CBP/p300, a common binding partner found in the ER co-activator complex (Zhang et al., 1996). Though STAT1 and ER did not directly bind to each other, these two factors could be connected by CBP/p300 or some other cofactor or combination of cofactors. In fact, STAT1 and ER are both present in the MegaTrans complex, a complex present at the most potent functional enhancers required for activation of enhancer RNA transcription and recruitment of coactivators (Liu et al., 2014). In addition, STAT3, another TF of the same family, has been shown to co-opt a subset of shared ER-FOXA-STAT3 enhancers to promote breast cancer metastasis independent of ER and FOXA1 through IL6/STAT3 signaling under non-estrogen withdrawn conditions (Siersbæk et al., 2020). In a similar manner, STAT1 could also be altering ER binding through its own INF/STAT1 signaling that is active particularly under low estrogenic conditions, which warrants further exploration.

STAT1 motif is also present with 10^−8^ M E2. Interestingly, more STAT1 motif fractions are found in Cluster 3 than with Clusters 4 and 5, suggesting STAT1 has a distinctive role in ER binding that predominates at lower E2 concentrations compared to higher E2. With enough E2/ER saturation to outcompete STAT1’s involvement due to higher E2 availability, the increased ER signaling may no longer need STAT1’s assistance in ER binding. STAT1, potentially as a coregulator, could influence the transcriptional potential of ER, allowing the fine-tuning of target gene transcription in response to the availability of E2 (Brisken & O’Malley, 2010). Future studies in conjunction with the other motifs we discovered, including STAT2, CPEB1, FOX family, and IRF family of transcription factors, some of which are related to STAT1, could illuminate the factors/complex involved in this process.

Further research is needed to understand the exact mechanism by which STAT1 influences ER binding, but previous reports validate STAT1’s involvement in breast cancer. Biopsies of human breast cancer also indicate elevated STAT1 expression can be a predictive marker for resistance to chemotherapy and radiation (Weichselbaum et al., 2008). We find a lower survival with higher STAT1 expression in ER^+^ breast cancer and activation of both MUC1 and STAT1 pathway in breast cancer has been reported to confer a poorer prognosis for patients (Khodarev et al., 2010). In an RNA-seq study between MCF7 and its tamoxifen-resistant LCC2 cell lines, STAT1 is shown to be important for ER signaling (Hou et al., 2018). It not only facilitates ER transcription and stimulates breast cancer cell proliferation, but it is also recruited to the ER promoter region, potentially conferring tamoxifen resistance. Depletion or inhibition of STAT1 results in decreased ER protein levels. We see higher ER expression at the lowest concentration that gradually decreases with more E2 presence, which could possibly be attributed to STAT1’s involvement in regulating ER expression and signaling.

In summary, we hypothesize that in estrogen withdrawn conditions, DNA functions as an ER sponge where ER is spending time in a different “neighborhood” and ER binding is driven by cofactors, hence many of these sites lack EREs. Once enough E2 becomes available, not only does ER’s binding mode change, but also its total levels decrease significantly. This lower protein level might focus activated ER binding to its strongest binding sites which contain EREs. This finding is supported by other studies which revealed prerecruitment of ER to some sites prior to E2 treatment (Hewitt et al., 2012). We think this binding funnels down to the main ER sites as a means to potentially prevent too much ER activation in a feedback loop that is accompanied by ER degradation.

Our results also seem to point towards ER shuttling to the chromatin prior to homeostatic turnover under hormone-depleted conditions as found in the study by Guan et al. (Guan et al., 2019), where increased ER in the chromatin fraction is present even with the absence of ER ligand with MG132 treatment. We detect nuclear ER immunofluorescence staining in all our experimental conditions including the negative control ETOH. Our study reveals the highest activity might be specific to the lower E2 states, representative of the most common, postmenopausal hormone receptor positive breast cancer status. Along with previous studies of STAT1 in ER^+^ breast cancer, we see the impact of STAT1 on ER binding that potentially dictates the breast cancer progression and treatment resistance reported. Thus, ER-STAT1 axis under the low E2 conditions could be a target to help improve the efficacy of the current anti-endocrine treatment modalities and halt tumor growth and the development of resistance. Our study’s framework of identifying non-ER motifs at low-E2 ER binding could be applied to find other factors that modulate ER binding at low E2, potentially leading to other ER-driven mechanisms at play in post-menopausal women.

## Acknowledgements

This study was supported by the NIH grant R01CA205044 (PK). We would like to thank the University of Colorado Cancer Center Genomics Shared Resource in sequencing our CUT&RUN and RNA-sequencing libraries (Cancer Center Support Grant P30CA046934), the laboratory of CAS for providing assistance with the Protein Simple-Jess Western Blot system.

## Declaration of interests

The authors declare there are no conflict of interests.

**Supplemental Figure 1. Immunofluorescence quantification of ER staining from the insets (zoomed 200% of 20X image) by Fiji (Fiji Is Just Image J)**.

**Supplemental Figure 2. UCD12 ER CUT&RUN**. A) Profile plots for spike-in normalized read density in different E2 concentrations indicating ER activity in UCD12 cells. B) Boxplot showing the CUT&RUN score distribution of the different E2/ETOH groups

**Supplemental Figure 3. ChromHMM-18 state model of MCF7 cells treated with 100nM E2 for 1 hour within the CUT&RUN clusters**.

**Supplemental Table 1. E2-peak stats**. Number of ERE vs. non-ERE peaks found in the specific clusters, distribution between proximal and distal.

## References

Alarid, E. T., Bakopoulos, N., & Solodin, N. (1999). Proteasome-mediated proteolysis of estrogen receptor: a novel component in autologous down-regulation. Mol Endocrinol, 13(9), 1522–1534. https://doi.org/10.1210/mend.13.9.0337

AlFakeeh, A., & Brezden-Masley, C. (2018). Overcoming endocrine resistance in hormone receptor-positive breast cancer. Curr Oncol, 25(Suppl 1), S18–s27. https://doi.org/10.3747/co.25.3752

Anurag, M., Ellis, M. J., & Haricharan, S. (2018). DNA damage repair defects as a new class of endocrine treatment resistance driver. Oncotarget, 9(91), 36252–36253. https://doi.org/10.18632/oncotarget.26363

Bailey, S. G., Cragg, M. S., & Townsend, P. A. (2012). Role of STAT1 in the breast. Jakstat, 1(3), 197–199. https://doi.org/10.4161/jkst.20967

Brechbuhl, H. M., Finlay-Schultz, J., Yamamoto, T. M., Gillen, A. E., Cittelly, D. M., Tan, A. C., Sams, S. B., Pillai, M. M., Elias, A. D., Robinson, W. A., Sartorius, C. A., & Kabos, P. (2017). Fibroblast Subtypes Regulate Responsiveness of Luminal Breast Cancer to Estrogen. Clin Cancer Res, 23(7), 1710–1721. https://doi.org/10.1158/1078-0432.Ccr-15-2851

Brisken, C., & O’Malley, B. (2010). Hormone action in the mammary gland. Cold Spring Harb Perspect Biol, 2(12), a003178. https://doi.org/10.1101/cshperspect.a003178

Bundred, N. J., Anderson, E., Nicholson, R. I., Dowsett, M., Dixon, M., & Robertson, J. F. (2002). Fulvestrant, an estrogen receptor downregulator, reduces cell turnover index more effectively than tamoxifen. Anticancer Res, 22(4), 2317–2319.

Carroll, J. S., Meyer, C. A., Song, J., Li, W., Geistlinger, T. R., Eeckhoute, J., Brodsky, A. S., Keeton, E. K., Fertuck, K. C., Hall, G. F., Wang, Q., Bekiranov, S., Sementchenko, V., Fox, E. A., Silver, P. A., Gingeras, T. R., Liu, X. S., & Brown, M. (2006). Genome-wide analysis of estrogen receptor binding sites. Nat Genet, 38(11), 1289–1297. https://doi.org/10.1038/ng1901

Cheung, E., & Kraus, W. L. (2010). Genomic analyses of hormone signaling and gene regulation. Annu Rev Physiol, 72, 191–218. https://doi.org/10.1146/annurev-physiol-021909-135840

Denver, N., Khan, S., Homer, N. Z. M., MacLean, M. R., & Andrew, R. (2019). Current strategies for quantification of estrogens in clinical research. J Steroid Biochem Mol Biol, 192, 105373. https://doi.org/10.1016/j.jsbmb.2019.04.022

Farcas, A. M., Nagarajan, S., Cosulich, S., & Carroll, J. S. (2021). Genome-Wide Estrogen Receptor Activity in Breast Cancer. Endocrinology, 162(2). https://doi.org/10.1210/endocr/bqaa224

Finlay-Schultz, J., Jacobsen, B. M., Riley, D., Paul, K. V., Turner, S., Ferreira-Gonzalez, A., Harrell, J. C., Kabos, P., & Sartorius, C. A. (2020). New generation breast cancer cell lines developed from patient-derived xenografts. Breast Cancer Res, 22(1), 68. https://doi.org/10.1186/s13058-020-01300-y

Fisher, B., Costantino, J. P., Wickerham, D. L., Redmond, C. K., Kavanah, M., Cronin, W. M., Vogel, V., Robidoux, A., Dimitrov, N., Atkins, J., Daly, M., Wieand, S., Tan-Chiu, E., Ford, L., & Wolmark, N. (1998). Tamoxifen for prevention of breast cancer: report of the National Surgical Adjuvant Breast and Bowel Project P-1 Study. J Natl Cancer Inst, 90(18), 1371–1388. https://doi.org/10.1093/jnci/90.18.1371

Gertz, J., Reddy, T. E., Varley, K. E., Garabedian, M. J., & Myers, R. M. (2012). Genistein and bisphenol A exposure cause estrogen receptor 1 to bind thousands of sites in a cell type-specific manner. Genome Res, 22(11), 2153–2162. https://doi.org/10.1101/gr.135681.111

Gilfillan, S., Fiorito, E., & Hurtado, A. (2012). Functional genomic methods to study estrogen receptor activity. J Mammary Gland Biol Neoplasia, 17(2), 147–153. https://doi.org/10.1007/s10911-012-9254-4

Guan, J., Zhou, W., Hafner, M., Blake, R. A., Chalouni, C., Chen, I. P., De Bruyn, T., Giltnane, J. M., Hartman, S. J., Heidersbach, A., Houtman, R., Ingalla, E., Kategaya, L., Kleinheinz, T., Li, J., Martin, S. E., Modrusan, Z., Nannini, M., Oeh, J., … Metcalfe, C. (2019). Therapeutic Ligands Antagonize Estrogen Receptor Function by Impairing Its Mobility. Cell, 178(4), 949-963.e918. https://doi.org/10.1016/j.cell.2019.06.026

Hah, N., Murakami, S., Nagari, A., Danko, C. G., & Kraus, W. L. (2013). Enhancer transcripts mark active estrogen receptor binding sites. Genome Res, 23(8), 1210–1223. https://doi.org/10.1101/gr.152306.112

Hewitt, S. C., Li, L., Grimm, S. A., Chen, Y., Liu, L., Li, Y., Bushel, P. R., Fargo, D., & Korach, K. S. (2012). Research resource: whole-genome estrogen receptor α binding in mouse uterine tissue revealed by ChIP-seq. Mol Endocrinol, 26(5), 887–898. https://doi.org/10.1210/me.2011-1311

Hou, Y., Li, X., Li, Q., Xu, J., Yang, H., Xue, M., Niu, G., Zhuo, S., Mu, K., Wu, G., Li, X., Wang, H., Zhu, J., & Zhuang, T. (2018). STAT1 facilitates oestrogen receptor α transcription and stimulates breast cancer cell proliferation. J Cell Mol Med, 22(12), 6077–6086. https://doi.org/10.1111/jcmm.13882

Keski-Rahkonen, P., Huhtinen, K., Desai, R., Harwood, D. T., Handelsman, D. J., Poutanen, M., & Auriola, S. (2013). LC-MS analysis of estradiol in human serum and endometrial tissue: Comparison of electrospray ionization, atmospheric pressure chemical ionization and atmospheric pressure photoionization. J Mass Spectrom, 48(9), 1050–1058. https://doi.org/10.1002/jms.3252

Khodarev, N., Ahmad, R., Rajabi, H., Pitroda, S., Kufe, T., McClary, C., Joshi, M. D., MacDermed, D., Weichselbaum, R., & Kufe, D. (2010). Cooperativity of the MUC1 oncoprotein and STAT1 pathway in poor prognosis human breast cancer. Oncogene, 29(6), 920–929. https://doi.org/10.1038/onc.2009.391

Kim, C., Tang, G., Pogue-Geile, K. L., Costantino, J. P., Baehner, F. L., Baker, J., Cronin, M. T., Watson, D., Shak, S., Bohn, O. L., Fumagalli, D., Taniyama, Y., Lee, A., Reilly, M. L., Vogel, V. G., McCaskill-Stevens, W., Ford, L. G., Geyer, C. E., Jr., Wickerham, D. L., … Paik, S. (2011). Estrogen receptor (ESR1) mRNA expression and benefit from tamoxifen in the treatment and prevention of estrogen receptor-positive breast cancer. J Clin Oncol, 29(31), 4160–4167. https://doi.org/10.1200/jco.2010.32.9615

Kolendowski, B., Hassan, H., Krstic, M., Isovic, M., Thillainadesan, G., Chambers, A. F., Tuck, A. B., & Torchia, J. (2018). Genome-wide analysis reveals a role for TDG in estrogen receptor-mediated enhancer RNA transcription and 3-dimensional reorganization. Epigenetics Chromatin, 11(1), 5. https://doi.org/10.1186/s13072-018-0176-2

Lei, J. T., Anurag, M., Haricharan, S., Gou, X., & Ellis, M. J. (2019). Endocrine therapy resistance: new insights. Breast, 48 Suppl 1(Suppl 1), S26–S30. https://doi.org/10.1016/s0960-9776(19)31118-x

Lin, Z., Reierstad, S., Huang, C. C., & Bulun, S. E. (2007). Novel estrogen receptor-alpha binding sites and estradiol target genes identified by chromatin immunoprecipitation cloning in breast cancer. Cancer Res, 67(10), 5017–5024. https://doi.org/10.1158/0008-5472.Can-06-3696

Liu, Z., Merkurjev, D., Yang, F., Li, W., Oh, S., Friedman, M. J., Song, X., Zhang, F., Ma, Q., Ohgi, K. A., Krones, A., & Rosenfeld, M. G. (2014). Enhancer activation requires trans-recruitment of a mega transcription factor complex. Cell, 159(2), 358–373. https://doi.org/10.1016/j.cell.2014.08.027

McDonnell, D. P., & Wardell, S. E. (2010). The molecular mechanisms underlying the pharmacological actions of ER modulators: implications for new drug discovery in breast cancer. Curr Opin Pharmacol, 10(6), 620–628. https://doi.org/10.1016/j.coph.2010.09.007

Mottamal, M., Kang, B., Peng, X., & Wang, G. (2021). From Pure Antagonists to Pure Degraders of the Estrogen Receptor: Evolving Strategies for the Same Target. ACS Omega, 6(14), 9334–9343. https://doi.org/10.1021/acsomega.0c06362

Nawaz, Z., Lonard, D. M., Dennis, A. P., Smith, C. L., & O’Malley, B. W. (1999). Proteasome-dependent degradation of the human estrogen receptor. Proc Natl Acad Sci U S A, 96(5), 1858–1862. https://doi.org/10.1073/pnas.96.5.1858

Osborne, C. K., & Schiff, R. (2011). Mechanisms of endocrine resistance in breast cancer. Annu Rev Med, 62, 233–247. https://doi.org/10.1146/annurev-med-070909-182917

Palaniappan, M., Nguyen, L., Grimm, S. L., Xi, Y., Xia, Z., Li, W., & Coarfa, C. (2019). The genomic landscape of estrogen receptor α binding sites in mouse mammary gland. PLoS One, 14(8), e0220311. https://doi.org/10.1371/journal.pone.0220311

Priyanka, H. P., Krishnan, H. C., Singh, R. V., Hima, L., & Thyagarajan, S. (2013). Estrogen modulates in vitro T cell responses in a concentration-and receptor-dependent manner: effects on intracellular molecular targets and antioxidant enzymes. Mol Immunol, 56(4), 328–339. https://doi.org/10.1016/j.molimm.2013.05.226

Rosner, W., Hankinson, S. E., Sluss, P. M., Vesper, H. W., & Wierman, M. E. (2013). Challenges to the measurement of estradiol: an endocrine society position statement. J Clin Endocrinol Metab, 98(4), 1376–1387. https://doi.org/10.1210/jc.2012-3780

Ross-Innes, C. S., Stark, R., Holmes, K. A., Schmidt, D., Spyrou, C., Russell, R., Massie, C. E., Vowler, S. L., Eldridge, M., & Carroll, J. S. (2010). Cooperative interaction between retinoic acid receptor-alpha and estrogen receptor in breast cancer. Genes Dev, 24(2), 171–182. https://doi.org/10.1101/gad.552910

Segars, J. H., Marks, M. S., Hirschfeld, S., Driggers, P. H., Martinez, E., Grippo, J. F., Brown, M., Wahli, W., & Ozato, K. (1993). Inhibition of estrogen-responsive gene activation by the retinoid X receptor beta: evidence for multiple inhibitory pathways. Mol Cell Biol, 13(4), 2258–2268. https://doi.org/10.1128/mcb.13.4.2258-2268.1993

Shang, Y., Hu, X., DiRenzo, J., Lazar, M. A., & Brown, M. (2000). Cofactor dynamics and sufficiency in estrogen receptor-regulated transcription. Cell, 103(6), 843–852. https://doi.org/10.1016/s0092-8674(00)00188-4

Siegel, R. L., Miller, K. D., Fuchs, H. E., & Jemal, A. (2022). Cancer statistics, 2022. CA Cancer J Clin, 72(1), 7–33. https://doi.org/10.3322/caac.21708

Siersbæk, R., Scabia, V., Nagarajan, S., Chernukhin, I., Papachristou, E. K., Broome, R., Johnston, S. J., Joosten, S. E. P., Green, A. R., Kumar, S., Jones, J., Omarjee, S., Alvarez-Fernandez, R., Glont, S., Aitken, S. J., Kishore, K., Cheeseman, D., Rakha, E. A., D’Santos, C., … Carroll, J. S. (2020). IL6/STAT3 Signaling Hijacks Estrogen Receptor α Enhancers to Drive Breast Cancer Metastasis. Cancer Cell, 38(3), 412-423.e419. https://doi.org/10.1016/j.ccell.2020.06.007

Skene, P. J., Henikoff, J. G., & Henikoff, S. (2018). Targeted in situ genome-wide profiling with high efficiency for low cell numbers. Nat Protoc, 13(5), 1006–1019. https://doi.org/10.1038/nprot.2018.015

Weichselbaum, R. R., Ishwaran, H., Yoon, T., Nuyten, D. S., Baker, S. W., Khodarev, N., Su, A. W., Shaikh, A. Y., Roach, P., Kreike, B., Roizman, B., Bergh, J., Pawitan, Y., van de Vijver, M. J., & Minn, A. J. (2008). An interferon-related gene signature for DNA damage resistance is a predictive marker for chemotherapy and radiation for breast cancer. Proc Natl Acad Sci U S A, 105(47), 18490–18495. https://doi.org/10.1073/pnas.0809242105

Yager, J. D., & Davidson, N. E. (2006). Estrogen carcinogenesis in breast cancer. N Engl J Med, 354(3), 270–282. https://doi.org/10.1056/NEJMra050776

Yaghjyan, L., & Colditz, G. A. (2011). Estrogens in the breast tissue: a systematic review. Cancer Causes Control, 22(4), 529–540. https://doi.org/10.1007/s10552-011-9729-4

Zelnak, A. B., & O’Regan, R. M. (2015). Optimizing Endocrine Therapy for Breast Cancer. J Natl Compr Canc Netw, 13(8), e56–64. https://doi.org/10.6004/jnccn.2015.0125

Zhang, J. J., Vinkemeier, U., Gu, W., Chakravarti, D., Horvath, C. M., & Darnell, J. E., Jr. (1996). Two contact regions between Stat1 and CBP/p300 in interferon gamma signaling. Proc Natl Acad Sci U S A, 93(26), 15092–15096. https://doi.org/10.1073/pnas.93.26.15092

