## Supplementary figures and images for "Estradiol (E2) concentration shapes the chromatin binding landscape of the estrogen receptor"

### Supplemental Figure 1

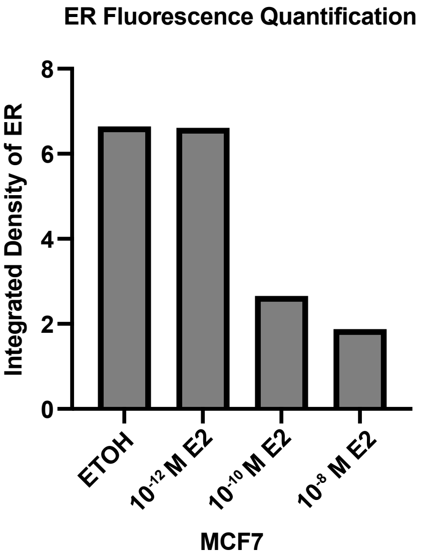

### Supplemental Figure 2

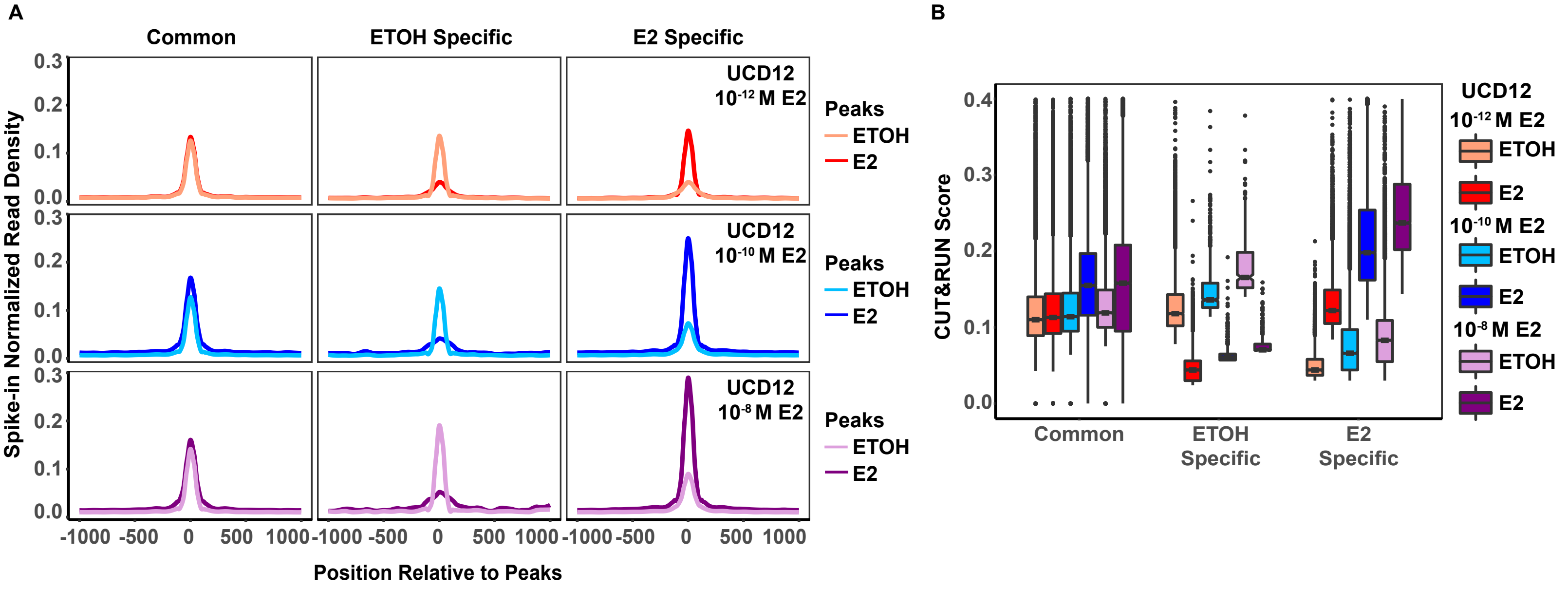

### Supplemental Figure 3

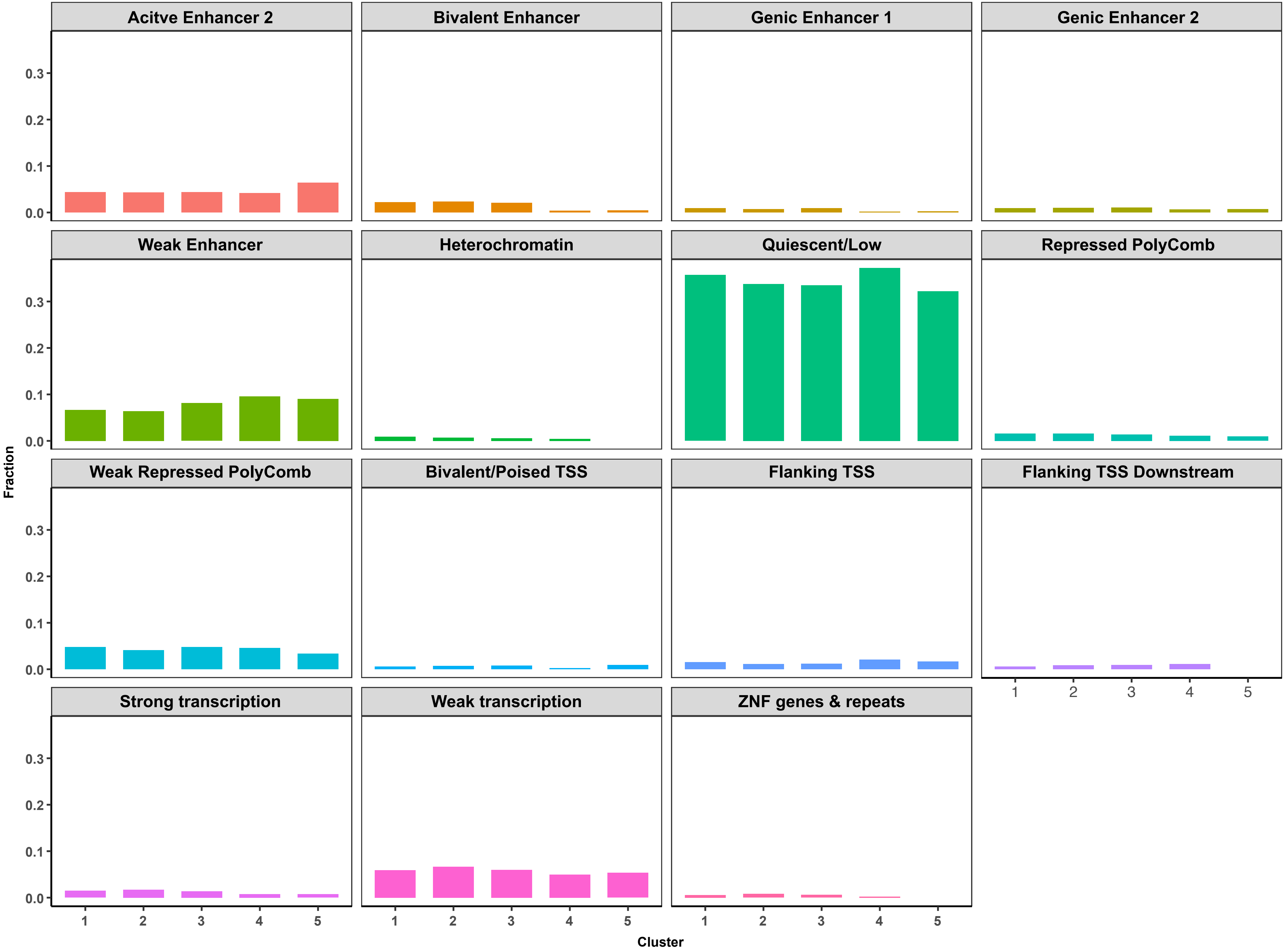
